# Evolutionary plasticity in the innate immune function of Akirin

**DOI:** 10.1101/230102

**Authors:** Jolanta Polanowska, Jia-Xuan Chen, Julien Soulé, Shizue Omi, Jerome Belougne, Clara Taffoni, Nathalie Pujol, Matthias Selbach, Olivier Zugasti, Jonathan J. Ewbank

## Abstract

Eukaryotic gene expression requires the coordinated action of transcription factors, chromatin remodelling complexes and RNA polymerase. The conserved nuclear protein Akirin plays a central role in immune gene expression in insects and mammals, linking the SWI/SNF chromatin-remodelling complex with the transcription factor NFκB. Although nematodes lack NFκB, Akirin is also indispensable for the expression of defence genes in the epidermis of *Caenorhabditis elegans* following natural fungal infection. Through a combination of reverse genetics and biochemistry, we discovered that in *C. elegans* Akirin has conserved its role of bridging chromatin-remodellers and transcription factors, but that the identity of its functional partners is different since it forms a physical complex with NuRD proteins and the POU-class transcription factor CEH-18. In addition to providing a substantial step forward in our understanding of innate immune gene regulation in *C. elegans*, our results give insight into the molecular evolution of lineage-specific signalling pathways.

## INTRODUCTION

A fundamental part of innate immune responses is the regulated expression of defence genes. In both vertebrates and many invertebrates, including *Drosophila*, two of the key regulators controlling innate immunity are the Rel-homology domain (RHD) protein NF-κB and its protein partner IκB (Lemaitre & Hoffmann, 2007). Across many species, NF-κB functions in concert with members of the conserved Akirin family (InterPro: IPR024132) to govern the expression of defence genes (Goto, Matsushita et al., 2008). More specifically, in vertebrates, Akirin 2 bridges NF-κB and the SWI/SNF chromatin-remodelling complex, by interacting with IκB-ζ and the BRG1-Associated Factor 60 (BAF60) proteins, downstream of Toll-like receptor (TLR) signalling (Tartey, Matsushita et al., 2014, Tartey & Takeuchi, 2015). In insects, an equivalent complex (including Relish and the Brahma-associated proteins BAP55 and BAP60 in *Drosophila*) governs the expression of antimicrobial peptide (AMP) gene expression upon infection by Gram-negative bacteria (Bonnay, Nguyen et al., 2014, Goto, Fukuyama et al., 2014, Tartey & Takeuchi, 2015).

Infection of *Caenorhabditis elegans* by its natural pathogen *Drechmeria coniospora* (Lebrigand, He et al., 2016) provokes an increase of AMP expression, but in the absence NF-κB and independently of the single TLR gene *tol-1* (Couillault, Pujol et al., 2004, Pujol, Link et al., 2001). It was therefore surprising that *akir-1*, the sole nematode Akirin orthologue was identified in a genome-wide RNAi screen for genes involved in the regulation of *nlp-29* (Squiban, Belougne et al., 2012, Zugasti, Thakur et al., 2016), an AMP gene that has been extensively used as a read-out of the epidermal innate immune response (e.g. (Couillault, Fourquet et al., 2012, Labed, Omi et al., 2012, Lee, Kniazeva et al., 2010, Tong, Lynn et al., 2009)).

These previous studies have revealed surprising molecular innovation in the pathways that regulate AMP gene expression. To give one example, in other animal species, STAT-like transcription factors function in concert with Janus kinases (JAKs). But in *C. elegans*, although there are no JAKs (Yamaoka, Saharinen et al., 2004), the 2 STAT-like proteins, STA-1 and STA-2, function in antiviral (Tanguy, Veron et al., 2017) and antifungal immunity (Dierking, Polanowska et al., 2011), respectively. In the latter case, STA-2’s function appears to be modulated by a nematode-specific member of the SLC6 family, SNF-12, acting downstream of the GPCR DCAR-1 and a p38 MAPK pathway to regulate *nlp-29* expression (Zugasti, Bose et al., 2014). Here, we undertook a focused study of *akir-1*, to understand how AMP gene expression is governed and also to gain insight into the evolution of lineage-specific signalling pathways.

## RESULTS

### The *C. elegans* Akirin homolog is required for antifungal innate immunity

We previously conducted a semi-automated genome-wide RNAi screen (Squiban et al., 2012) for genes that control the expression of the AMP reporter gene *nlp-29p::gfp*, following infection of *C. elegans* with *D. coniospora* (Zugasti et al., 2016). The candidates identified as positive regulators are collectively referred to as Nipi genes, for No Induction of Peptide expression after Infection. While *akir-1*(RNAi) caused a robust reduction in the induction of *nlp-29p::gfp* expression after infection (Fig 1A), it did not significantly affect the size of treated worms, nor the expression of the control *col-12p::DsRed* reporter transgene (Fig S1A), identifying it as Nipi gene and suggesting that it could have a specific function in innate immunity. When we used an available deletion allele, *akir-1*(*gk528*), which is predicted to be a molecular null, we recapitulated the effect on *nlp-29p::gfp* expression (Fig S1B). This analysis was, however, hampered by the mutants’ pleiotropic phenotypes (Clemons, Brockway et al., 2013), including a developmental delay and very marked decrease in the expression of the control reporter transgene (Fig S1C). To avoid these confounding effects, and since RNAi of *akir-1* gave robust and reproducible results, we used *akir-1*(RNAi) for our subsequent analyses.

**Figure 1.**
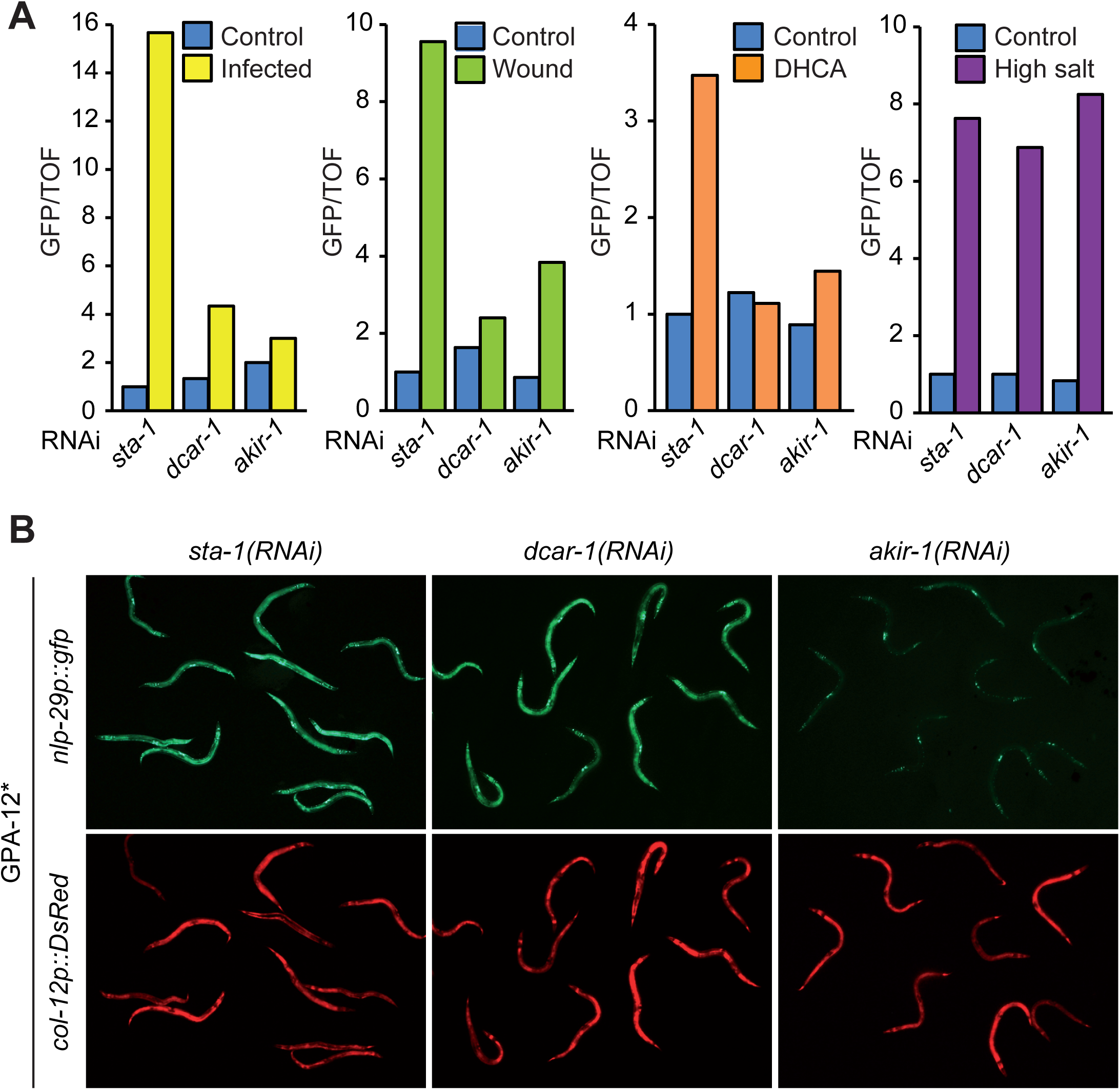
Akirin acts downstream of Gα to regulate the expression of *nlp-29*. **A.** Ratio of green fluorescence (GFP) to size (time of flight; TOF) in worms carrying the integrated array *frIs7* (containing *nlp-29p::gfp* and *col-12p::DsRed)* treated with RNAi against negative and positive controls (*sta-1, dcar-1*, respectively) or *akir-1* and infected by *D. coniospora* (Infected), wounded (Wound), treated with a 5 mM dihydrocaffeic acid solution (DHCA) or exposed to 300mM NaCl (High salt). Here and in subsequent figures representing Biosort results, unless otherwise stated, graphs are representative of at least 3 independent experiments with a minimum of 50 worms analysed for each condition. In this and subsequent figures, as previously explained (Pujol et al., 2008a), error bars are intentionally omitted. **B.** Fluorescent images of adult worms carrying *frIs7*, expressing a constitutively active Gα protein, GPA-12*, in the epidermis and treated with RNAi against the indicated genes. Almost all of the residual GFP expression seen upon *akir-1*(RNAi), most prominent in the vulval muscle cells, comes from *unc-53Bp::gfp* used as a transgenesis marker.

The induction of *nlp-29p::gfp* expression upon *D. coniospora* infection is correlated to the infectious burden, which in turn reflects the propensity of spores to bind the worm cuticle (Rouger, Bordet et al., 2014, Zugasti et al., 2016). There was no reduction in spore adhesion following *akir-1*(RNAi) (EV1D). Many genes required for the induction of *nlp-29p::gfp* expression after infection, including the GPCR gene *dcar-1* (Zugasti et al., 2014) and the STAT transcription factor-like gene *sta-2* (Dierking et al., 2011), are also required for the transcriptional response of *C. elegans* to physical injury. We found that *akir-1*(RNAi) also abrogated reporter transgene expression upon wounding (Fig 1A). One trigger for the epidermal innate immune response is the increase in the tyrosine metabolite HPLA that accompanies infection with *D. coniospora*. HPLA acts via DCAR-1 to activate a p38 MAPK signalling cascade (Zugasti et al., 2014). This GPCR can also be activated by the HPLA tautomer DHCA (Aoki, Yagami et al., 2011), a non-physiological ligand, which we use routinely as it is somewhat more potent and less toxic for worms than HPLA (Zugasti et al., 2014). The induction of *nlp-29p::gfp* expression upon exposure to DHCA was greatly reduced upon *akir-1*(RNAi), to a degree that was comparable to *dcar-1*(RNAi) (Fig 1A). Together, these results indicate that *akir-1* is required for the activation of the epidermal innate immune response, downstream of DCAR-1.

**Figure EV1.**
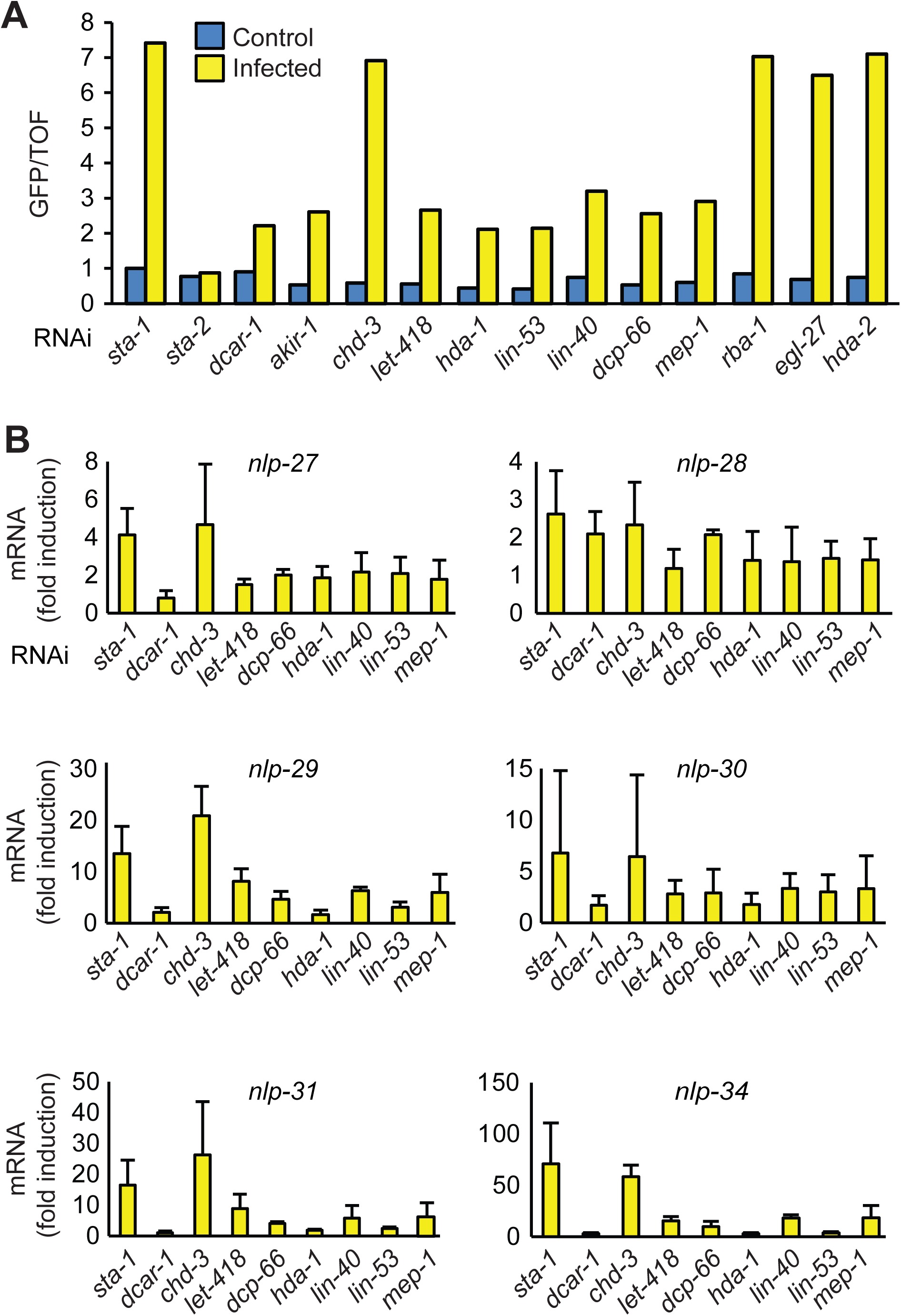
LET-418 NuRD and MEC complexes act in the epidermis to modulate *nlp* AMP gene expression. **A.** Ratio of green fluorescence (GFP) to size (TOF) in *rde-1*(*ne219*);*wrt-2p::RDE-1* worms that are resistant to RNAi except in the epidermis carrying the array *frIs7* treated with RNAi against the indicated genes and then infected or not by *D. coniospora*. **B.** Quantitative RT-PCR analysis of the fold induction of expression of genes in the *nlp-29* cluster in *rde-1*(*ne219*); *wrt-2p::RDE-1* worms treated with RNAi against the indicated genes, comparing expression levels in worms infected by *D. coniospora* with uninfected worms. The 6 RNAi clones block the induction of expression of each of the endogenous *nlp* AMP genes more or less completely, with the exception of *nlp-28*. Data are from three independent experiments (average and SD).

In contrast to the induction of *nlp-29p::gfp* provoked by infection, wounding or DHCA, the induction of *nlp-29p::gfp* observed after 6 hours exposure to moderate osmotic stress is DCAR-1 and p38 MAPK PMK-1-independent (Pujol, Zugasti et al., 2008b, Zugasti et al., 2014). We found that *akir-1*(RNAi), like *dcar-1*(RNAi), did not affect the induction of reporter gene expression upon osmotic stress (Fig 1A). Unlike *dcar-1*(*RNAi*), but similar to *sta-2(RNAi)* (Labed et al., 2012, Ziegler, Kurz et al., 2009), *akir-1(RNAi)* abolished the strong expression of *nlp-29p::gfp* seen in worms expressing a constitutively active form of the Gα protein GPA-12 (GPA-12*) (Fig 1B). Together, these results support the specific role for *akir-1* in innate immune signalling, placing it downstream of, or in parallel to, *gpa-12*.

#### *akir-1* acts in an epidermis-specific manner

To evaluate when and where *akir-1* was expressed, we generated strains carrying a transcriptional reporter gene (*akir-1p::gfp*). Consistent with previous studies (Hunt-Newbury, Viveiros et al., 2007), expression of GFP was observed from the late embryo stage onwards, peaking at the late L4 stage. Expression was most evident in the lateral epithelial seam cells, the major epidermal syncytium, hyp7, as well as in multiple head and tail neurons (Fig 2A). The different components of the p38 MAPK pathway, including *dcar-1, gpa-12* and *sta-2,* act in a cell autonomous fashion in the epidermis (Dierking et al., 2011, Labed et al., 2012, Zugasti et al., 2014). To determine whether this was also the case for *akir-1*, we knocked down its expression specifically in the epidermis. This greatly decreased *nlp-29p::gfp* expression upon infection, and also, as judged by qRT-PCR substantially reduced the induction of all the genes of the *nlp-29* locus, while not affecting their constitutive expression (Fig 2B,C & S2A). This indicates that *akir-1* acts in a cell autonomous manner in the epidermis to modulate AMP gene expression upon infection.

**Figure 2.**
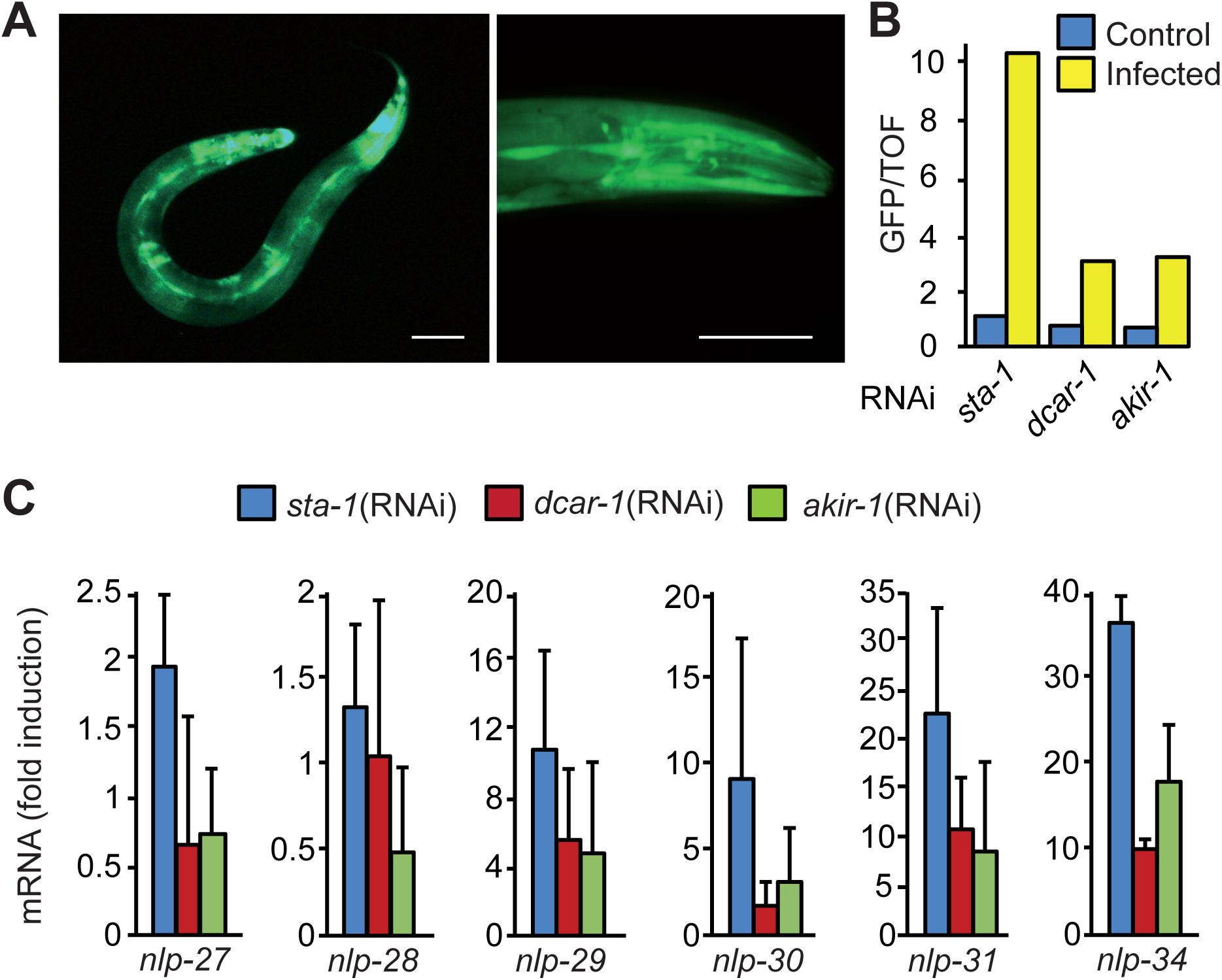
Akirin acts in the epidermis to regulate multiple *nlp* genes. **A.** Confocal images of IG1485 transgenic worms expressing an *akir-1p::gfp* reporter gene showing epidermal and neuronal expression of GFP. Scale bar 50 μm. **B.** Ratio of green fluorescence (GFP) to size (TOF) in *rde-1*(*ne219*);*wrt-2p::RDE-1* worms that are resistant to RNAi except in the epidermis carrying the array *frIs7* treated with RNAi against the indicated genes and infected by *D. coniospora.* C. Quantitative RT-PCR analysis of the expression of genes in the *nlp-29* cluster in *rde-1*(*ne219*); *wrt-2p::RDE-1* worms treated with RNAi against the indicated genes and infected by *D. coniospora*; results are presented relative to those of uninfected worms. Data are from three independent experiments (average and SD).

To test the functional relevance of these observations, we assayed the effect of epidermis-specific *akir-1*(RNAi) on the resistance of *C. elegans* to *D. coniospora* infection. Knocking down *akir-1* in this way was associated with a significant reduction in survival (Fig 3). Interpretation of this result is complicated by the fact that the same RNAi treatment also caused a significant decrease in longevity on non-pathogenic *E. coli* (Fig S2B), so the reduced resistance to *D. coniospora* infection is not likely to result solely from the observed diminution in AMP gene expression.

**Figure 3.**
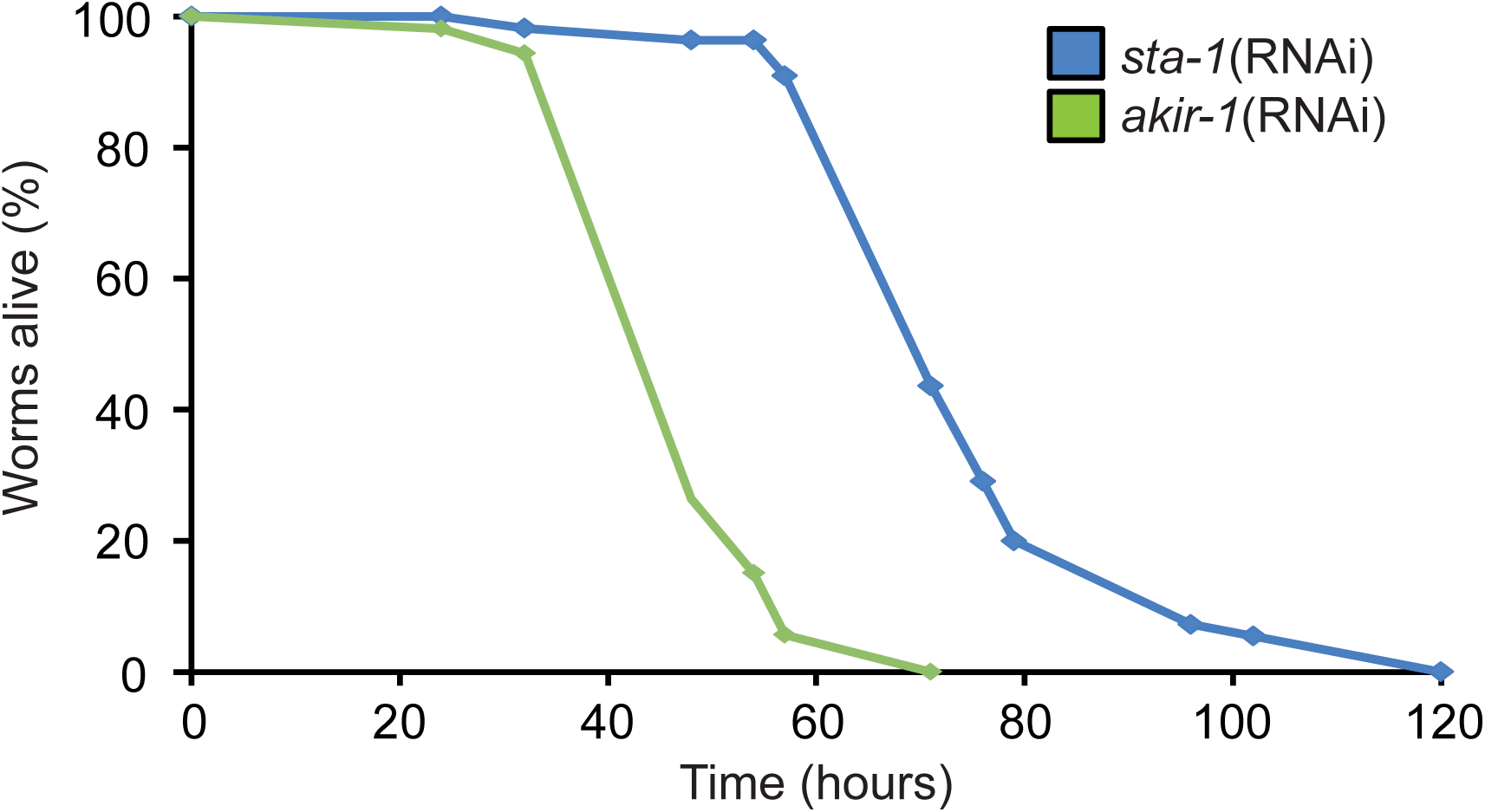
Akirin expression in the epidermis regulates resistance to fungal infection. Survival of *rde-1*(*ne219*);*wrt-2p::RDE-1* worms treated with RNAi against *sta-1* (n=50) or *akir-1* (n=50). The difference between the *sta-1(RNAi)* and *akir-1(RNAi)* animals is highly significant (p<0.0001; one-sided log rank test). Data are representative of three independent experiments.

#### *akir-1* and chromatin remodelling

In both mice and flies, Akirin controls innate immune gene expression through its interaction with BAF60/BAP60 and more generally the SWI/SNF chromatin-remodelling complex (Bonnay et al., 2014, Goto et al., 2008, Tartey et al., 2014). We therefore used RNAi to knock down the expression of components of the nematode SWI/SNF chromatin-remodelling complexes, but also of the Nucleosome Remodelling and histone Deacetylase (NuRD) and MEC complexes, as well as related genes (Passannante, Marti et al., 2010). With the exception of *swsn-1*, which caused pleiotropic development defects and affected expression of the control *col-12p::DsRed* reporter transgene, consistent with our previous results (Zugasti et al., 2016), none of the other SWI/SNF genes appeared to be required for *nlp-29p::gfp* expression (Fig S3A,B). On the other hand, knocking down 6 genes *dcp-66, hda-1, let-418, lin-40, lin-53*, and *mep-1*, largely, and specifically, blocked the expression of *nlp-29p::gfp* upon *D. coniospora* infection (Fig 4A and S3C,D). Of note, the 3 RNAi clones that gave the most robust Nipi phenotype, those targeting *hda-1/HDAC, lin-40/MTA* and *dcp-66/p66*, had been identified in the previous genome-wide screen (Zugasti et al., 2016). These 3 genes encode core subunits of the two canonical chromatin-remodelling (NuRD) complexes in *C. elegans*. The two complexes also share LIN-53/RbAp, but differ in their Mi-2 orthologs, having either LET-418 or CHD-3. LET-418 but not CHD-3, can interact with the Krüppel-like protein MEP-1 in a distinct complex, the MEC complex (Passannante et al., 2010, Xue, Wong et al., 1998). Our results suggest that both the LET-418-containing NuRD complex and the MEC complex are involved in defence gene expression. The 6 RNAi clones also strongly abrogated the elevated expression of *nlp-29p::gfp* normally seen in worms expressing GPA-12*, in clear contrast to *chd-3*(RNAi) (Fig 4B). Epidermis-specific RNAi with the same 6 clones also blocked the induction of reporter gene expression (Fig EV1A). Under these conditions, the induction of expression of 5 endogenous *nlp* AMP genes normally provoked by *D. coniospora* infection was also severely compromised (Fig EV1B). In contrast, there was no evidence for a role for HDA-2, RBA-1 or EGL-27 (Fig 4A), the respective homologues of the core subunits HDA-1, LIN-53 and LIN-40, that do not form part of either of the 2 biochemically characterized NuRD complexes in *C. elegans* (Passannante et al., 2010). Together these results suggest that both the LET-418 containing NuRD complex and the MEC complex act cell autonomously in the epidermis, downstream of (or in parallel to) *gpa-12*, to control *nlp* AMP gene expression upon *D. coniospora* infection. Further, they suggest that in contrast to what has been described in flies and mammals, AMP gene expression is not dependent upon the SWI/SNF complex in *C. elegans*, and raised the possibility that AKIR-1 might function together with the NuRD and MEC chromatin remodelling complexes.

**Figure 4.**
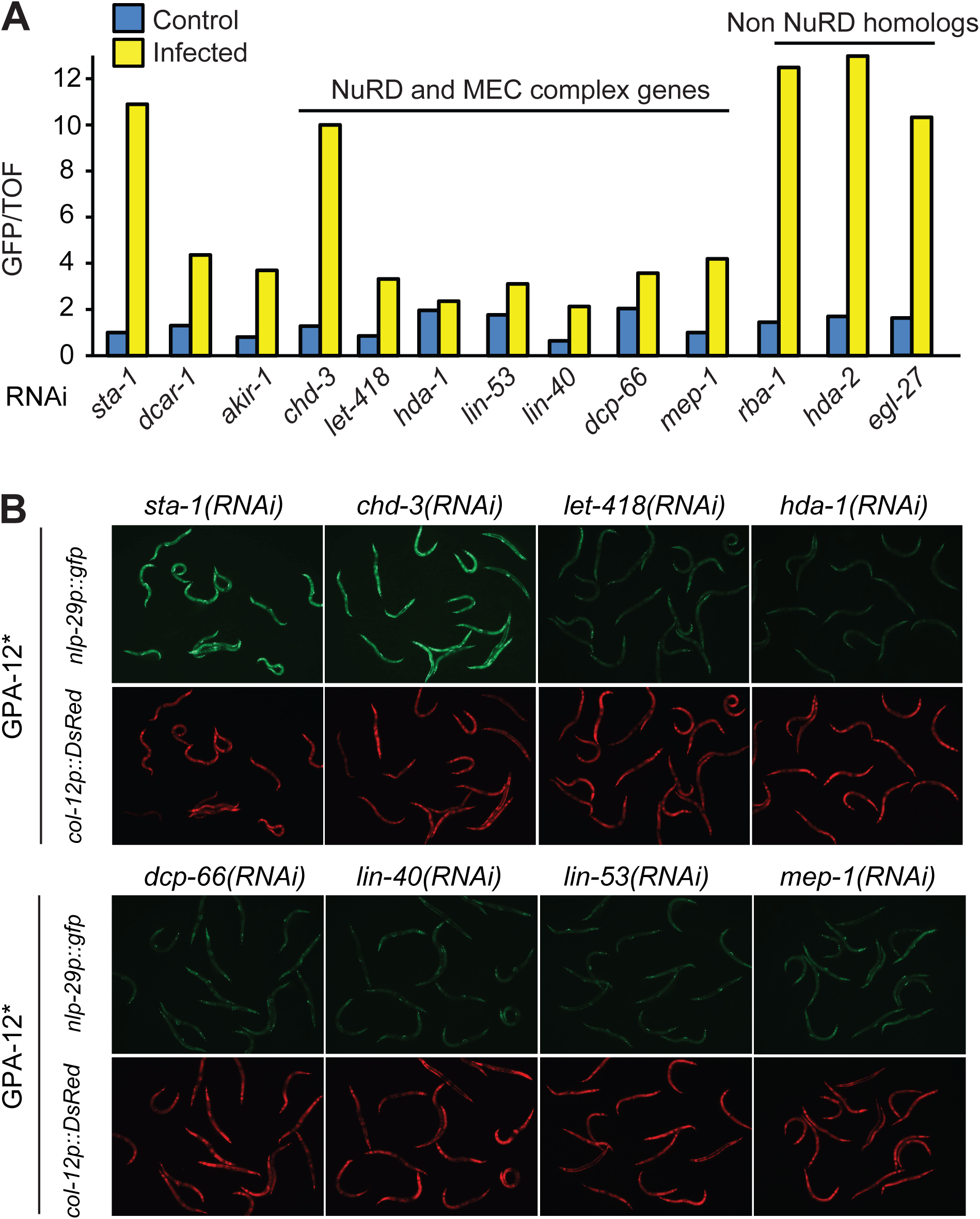
LET-418 NuRD and MEC complexes regulate *nlp-29* gene expression. **A.** Ratio of green fluorescence (GFP) to size (TOF) in worms carrying *frIs7,* treated with RNAi against control *(sta-1, dcar-1, akir-1)*, NuRD and MEC complex component, and non-NuRD chromatin remodelling component genes, and infected or not with *D. coniospora*. **B.** Fluorescent images of adult worms carrying *frIs7* and expressing GPA-12* in the epidermis, and treated with RNAi against the indicated genes. See legend to Figure 2 for more details.

### AKIR-1 forms a complex with components of the NuRD and MEC chromatin remodelling complexes and CEH-18

To address this possibility, we took an unbiased biochemical approach to identify the *in vivo* protein partners of AKIR-1. From a mixed-stage population of worms carrying a functional *akir-1p::AKIR-1::gfp* construct (Fig EV2), we pulled down AKIR-1::GFP by immunoprecipitation from whole worm extracts and subjected the purified proteins to mass spectrometry analysis (Figure 5A). Remarkably, all of the proteins that make up the NuRD and MEC complexes were found, i.e. LIN-40, LIN-53, LET-418, HDA-1, MEP-1 and DCP-66. The first 3, together with 6 other known or putative DNA-binding or transcription-related proteins (Reece-Hoyes, Deplancke et al., 2005), including CEH-18, were among the 53 high confidence protein partners (Figure 5B, C). Significantly, 9 of these 53 candidates (p=2.7x10^−7^), again including LIN-40 and CEH-18, had been identified in our previous RNAi screen for regulators of *nlp-29p::gfp* (Zugasti et al., 2016). In the complete list of close to 1400 protein partners, there were a further 111 hits (Table S1), so overall, fully 35% of the known candidate regulators of AMP gene expression (Nipi genes) were recovered through this independent biochemical approach when one includes the lower confidence candidates.

**Figure EV2.**
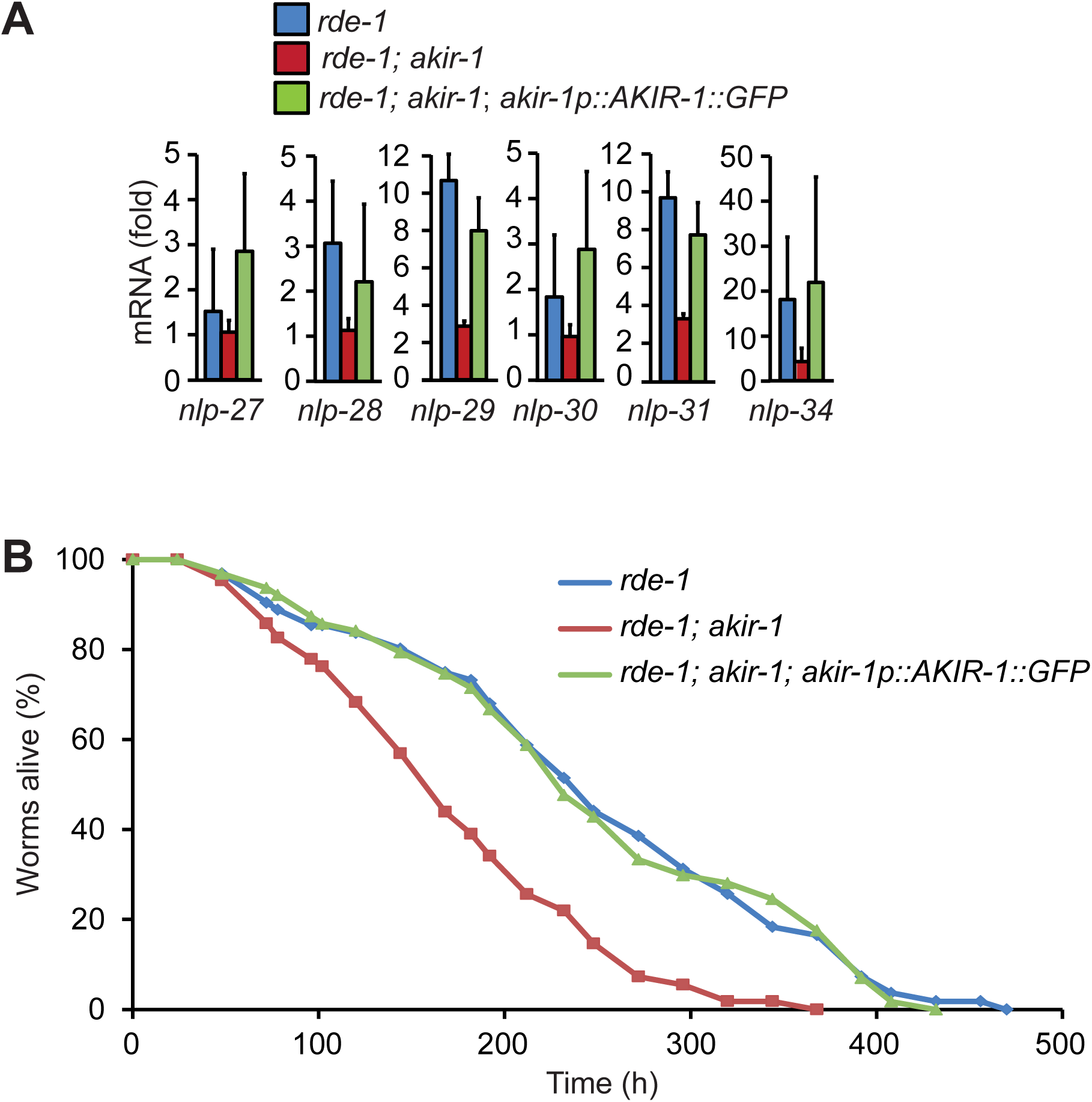
Transgenic rescue of *akir-1* mutant phenotypes. **A.** Quantitative RT-PCR analysis of the fold induction of expression of genes in the *nlp-29* cluster in *rde-1*(*ne219*), *rde-1*(*ne219*);*akir-1*(*gk528*) and *rde-1*(*ne219*);*akir-1*(*gk528*); *akir-1p::AKIR-1::gfp* worms, comparing expression levels in worms infected by *D. coniospora* with uninfected worms. Data are from three independent experiments (average and SD). **B.** Lifespan of *rde-1*(*ne219*), *rde-1*(*ne219*);*akir-1*(*gk528*) and *rde-1*(*ne219*);*akir-1*(*gk528*); *akir-1p::AKIR-1::gfp* worms. Data are representative of three independent experiments.

**Figure 5.**
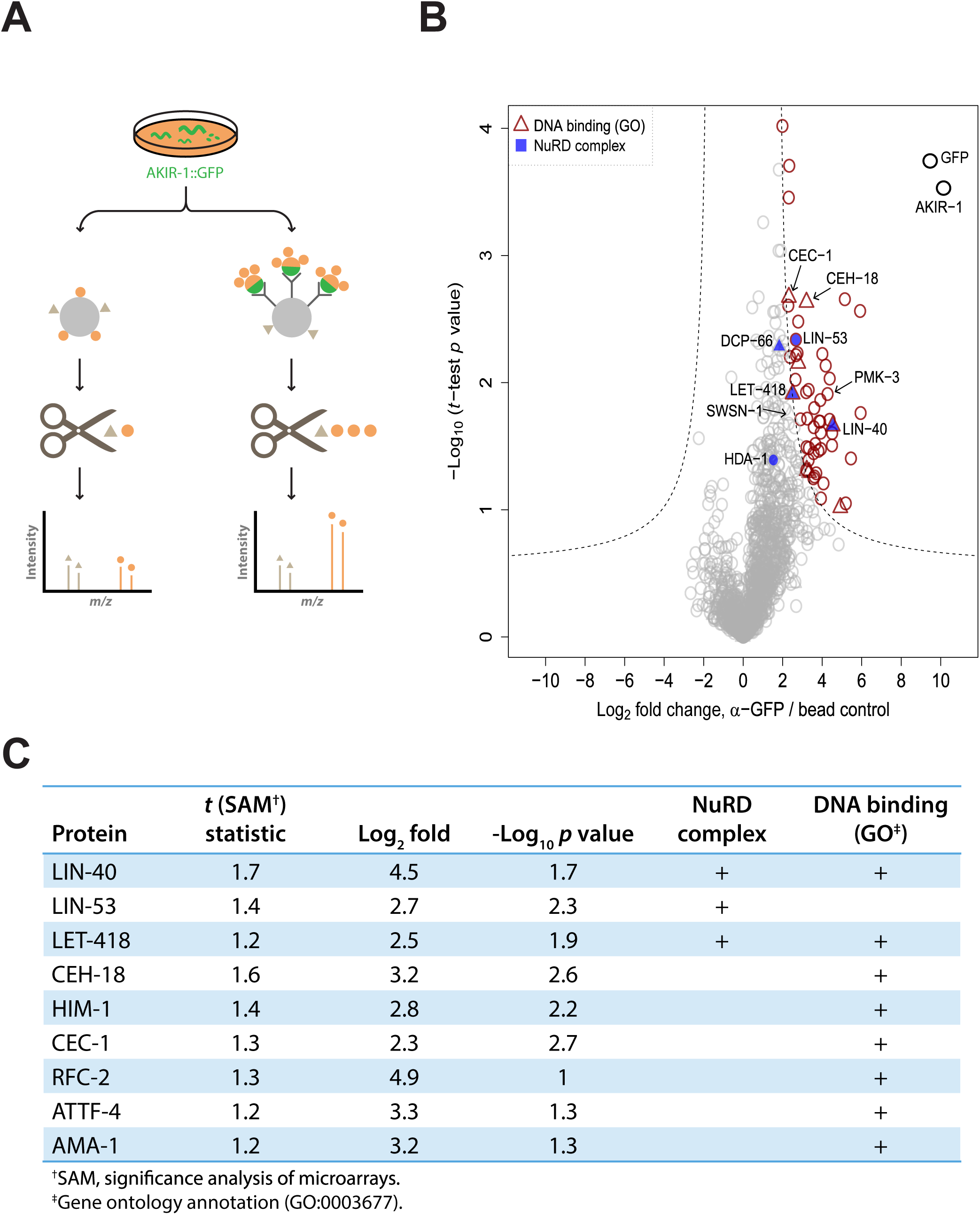
AKIR-1 interactors identified by label-free quantitative immunoprecipitation. **A.** Experimental design. Protein extracts from mixed-stage worms expressing AKIR-1::GFP were incubated with anti-GFP conjugated or control resins before proteolytic release of peptides from the immunoprecipitated proteins. The relative abundance of co-precipitated proteins was assessed by mass spectrometry. **B.** Volcano plot showing specific interaction partners (in red) of AKIR-1::GFP. The mean values for fold change from 3 independent experiments are shown. The SAM (significance analysis of microarrays) algorithm was used to evaluate the enrichment of the detected proteins. Proteins that met the combined enrichment threshold (hyperbolic curves, *t_0_* = 1.2) are colored in red. Proteins with the gene ontology annotation “DNA-binding” (GO:0003677) are depicted as triangles. Known members of the NuRD complex are shown in blue. C. NuRD complex and/or DNA-binding proteins among the 53 high confidence AKIR-1::GFP interaction partners.

When we compared the list of 53 high confidence candidate AKIR-1 binding proteins with the 190 proteins identified as potential interactors of the nematode BAP60 homologue SWSN-2.2 (Ertl, Porta-de-la-Riva et al., 2016), we found only 3 common proteins, none of which have been characterized as being specific regulators of *nlp-29p::gfp* expression (i.e. found as Nipi genes (Zugasti et al., 2016); Table S2). Using a less stringent list of 190 potential AKIR-1 binding proteins extended the overlap to 11 common partners, with just 2 corresponding to Nipi genes (*arp-1* and *dlst-1* that encode an actin-related protein, and a predicted dihydrolipoyllysine succinyltransferase, respectively). The 11 common proteins did, however, also include SWSN-1 and SWSN-4 (Table S2). This suggests that in some contexts, but not during its regulation of AMP gene expression, AKIR-1 might interact with the SWI/SNF complex. This functional dichotomy was further reinforced by examining the genes differentially regulated following knockdown of *swsn-2.2* and its paralogue *ham-3* (Ertl et al., 2016). There were only a very small number (28/1468) of genes characterized as up-regulated by *D. coniospora* infection and among them, there were none encoding AMPs (JJE unpublished observations). Together these results support the idea that there is a specific AKIR-1-containing protein complex involving the NuRD and MEC chromatin remodellers, required for AMP gene regulation.

We therefore focused on the interaction between AKIR-1 and these chromatin-remodelling factors. We used available antibodies to validate the NuRD and MEC complex proteins LET-418 and HDA-1 as AKIR-1-interactors. Both could be detected together with AKIR-1::GFP, in samples from infected and control worms, derived the strain used for mass spectrometric analysis, and importantly also from a strain of worms carrying a single copy *akir-1::gfp* insertion in the wild-type background (Fig 6A, EV3). There was a clear reduction in the quantity of LET-418 that was pulled down with AKIR-1::GFP from the samples of infected worms compared to non-infected worms. The same tendency was observed for HDA-1. These results strongly support the existence of a physical complex between AKIR-1 and the NuRD and MEC complexes in uninfected worms that changes following infection.

**Figure 6.**
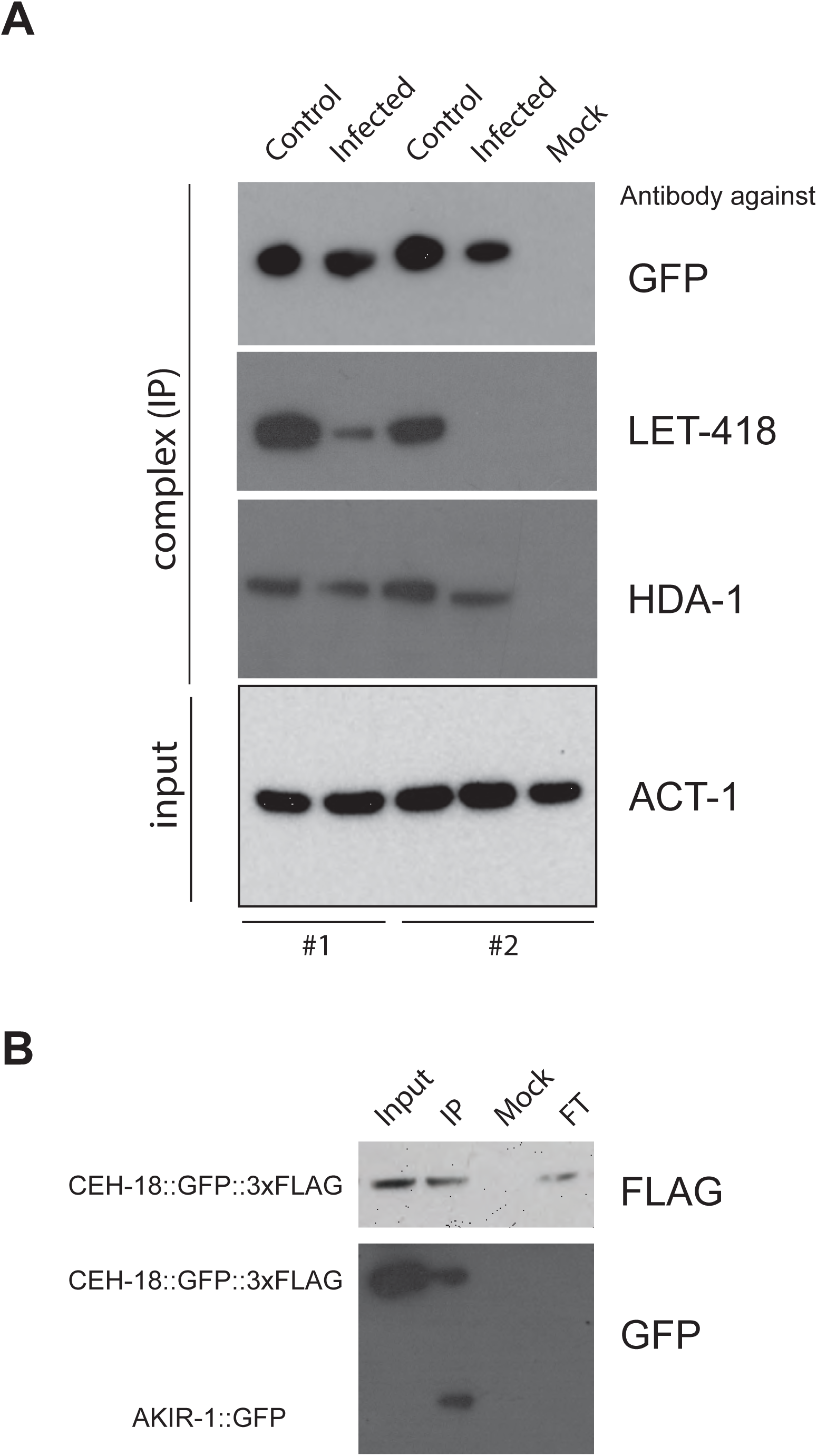
Validation of AKIR-1 interactors by Western blotting. **A.** Complexes immunopurified using an anti-GFP antibody from control or infected worms with a single copy *AKIR-1::GFP* insertion (wt; *frSi12[pNP157(akir-1p::AKIR-1::GFP)] II*) were probed with specific antibodies. The results for two independent pull-downs are shown. The presence of HDA-1 and LET-418 (NuRD complex components) could be confirmed. Anti-ACT-1 was used to control the total input for each immunoprecipitation. **B.** Complexes immunopurified using an anti-FLAG antibody, from a strain co-expressing AKIR-1::GFP and FLAG-tagged CEH-18 (*wt; frSi12[pNP157(akir-1p::AKIR-1::GFP)] II; wgIs533[CEH-18::TY1::GFP::3xFLAG + unc-119(+)]*), were probed with anti-FLAG (top panel) and anti-GFP (bottom) antibodies. In addition to the immunopurified complex (IP), the extract before immunopurification (Input), the unbound fraction (flow-through: FT) and proteins immunopurified using an unrelated antibody (Mock) were also analysed.

**Figure EV3.**
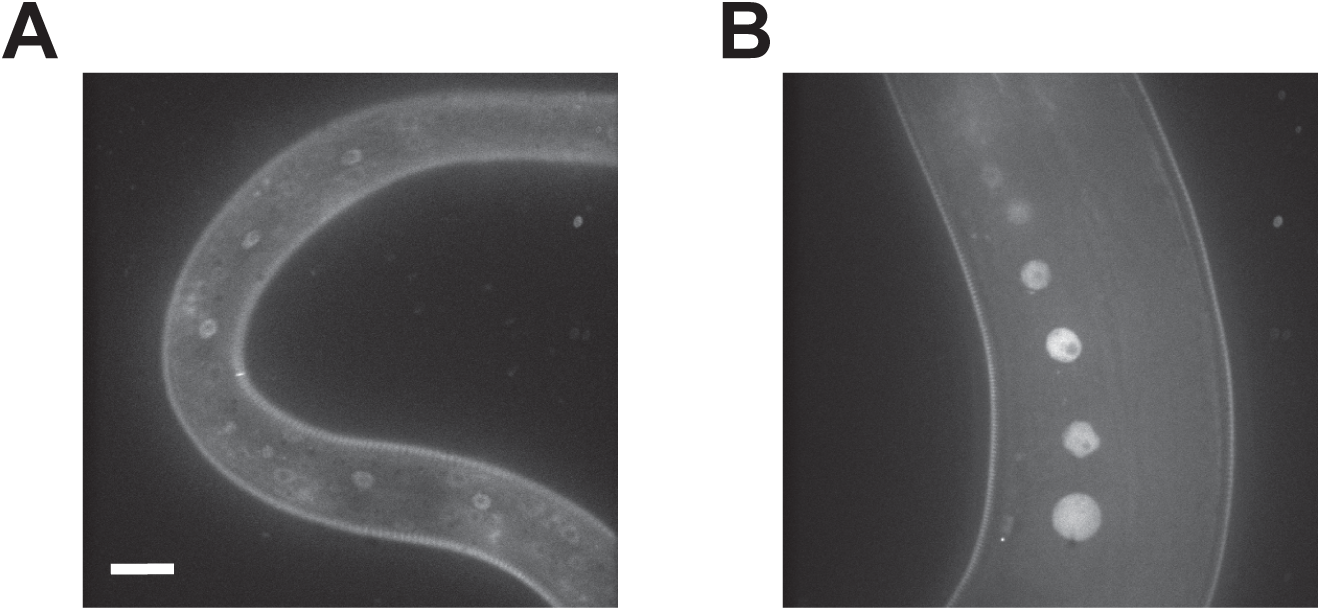
Expression pattern of AKIR-1::GFP. Worms carrying a single copy insertion of an AKIR-1::GFP construct (*wt; frSi12[pNP157(akir-1p::AKIR-1::GFP)] II*) were visualized by confocal microscopy and several confocal planes summed. AKIR-1::GFP showed a clear and strong nuclear localization: **A.** epidermal nuclei in an L3 stage worm **B.** Germline nuclei in a young adult. The scale bar is 20 μm.

When we analysed the complex that was pulled-down together with CEH-18, using a strain expressing a doubly-tagged version of the protein (CEH-18::GFP::3xFLAG; (Zhong, Niu et al., 2010)), we were readily able to detect AKIR-1 (Fig 6B), lending further support to the proposed AKIR-1/NuRD/CEH-18 complex.

### AKIR-1 binds to AMP gene promoters

We then addressed the question of whether AKIR-1 has the potential to interact with DNA, by chromatin immunoprecipitation (ChIP), using the strain of worms carrying a single copy *akir-1::gfp* insertion. We assayed its capacity to associate with DNA fragments corresponding to the promoters of 3 AMP genes, or to their 3’ UTRs, using samples from uninfected or infected populations of worms. For all 3 AMP genes assayed, we detected DNA binding and observed markedly preferential binding to the promoter regions relative to the 3’ UTRs. The 3 genes, *nlp-29, nlp-31* and *nlp-34* are strongly induced by *D. coniospora* infection (Pujol et al., 2008b). We observed a more than 10-fold higher occupancy of AKIR-1 on DNA in the samples from non-infected worms relative to the infected ones (Figure 7A). Taken together, our results support a model, discussed further below, in which AKIR-1 plays 2 indissociable roles. First, in association with the NuRD and MEC complexes, it binds to the promoters of defence genes and potentially recruits transcription factors including CEH-18. Secondly, AKIR-1 and its protein partners negatively regulate the transcription of defence genes, with this repression being relieved upon their removal from their binding sites following infection (Figure 7B). This could explain why loss of AKIR-1 function is associated with incapacity to express AMP genes upon infection.

**Figure 7.**
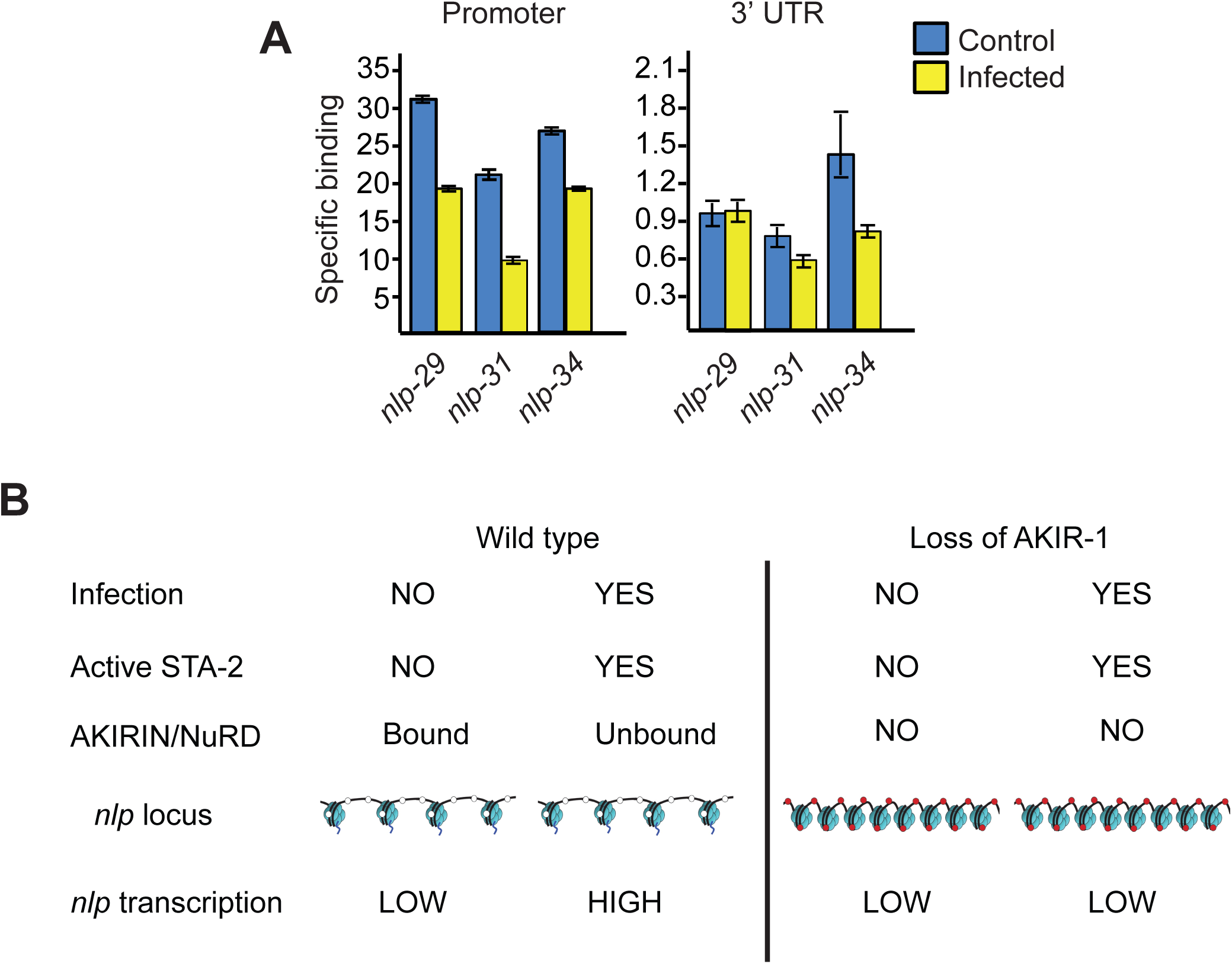
AKIR-1 binds preferentially to *nlp* gene promoters in the absence of infection. **A.** Specific binding of AKIR-1::GFP onto promoters (left panel) or 3’ UTR (right panel) of *nlp-29, nlp-31* and *nlp-34*, represented as the fold enrichment of the specific ChIP signal obtained using an anti-GFP antibody for immunoprecipitation relative to that when blocked beads were used, measured by quantitative PCR. Data is normalised to input; the average (and standard error) from three independent experiments is shown. **B.** Model for the role of AKIR-1 in the regulation of *nlp* AMP gene expression upon infection. The image of DNA/chromatin is adapted from P. Luong, Basic Principles of Genetics (cnx.org) under a Creative Commons Attribution License.

## DISCUSSION

We are interested in the mechanisms involved in the regulated expression of *nlp-29*, a representative of one class of AMP genes in *C. elegans* (Pujol, Cypowyj et al., 2008a, Pujol, Davis et al., 2012, Pujol et al., 2008b). In common with many other AMP genes, the level of *nlp-29* mRNA rapidly increases following either physical injury or infection with the nematophagous fungus *D. coniospora.* In both cases, the integrity of the cuticle and underlying epidermis is compromised. Although we have advanced in our understanding of how this triggers the innate immune response, and how the associated signal transduction pathway is organized, the details of the transcriptional regulation remain to be fully elucidated. We previously identified ELT-3, an epidermis specific GATA factor as being partially required, in a generic fashion, for *nlp-29* expression (Pujol et al., 2008b). The STAT-like transcription factor STA-2 plays a more specific role. It is largely dispensable for the constitutive expression of *nlp-29*, but is required for its induction upon wounding and infection (Dierking et al., 2011, Labed et al., 2012). In this work, we have made a considerable step forward by characterizing the key role of AKIR-1 and identifying its protein partners, including the NuRD and MEC complex chromatin remodelling proteins and the transcription factor CEH-18. All these factors are required for AMP gene expression after fungal infection of the nematode epidermis.

CEH-18 is a member of the POU subgroup of the Hox class of homeodomain transcription factors. These are regulators of cellular proliferation, differentiation and migration across species. In *C. elegans, ceh-18* has primarily been characterized for its negative regulatory role in a somatic gonadal sheath cell-dependent pathway that governs oocyte meiotic arrest (Miller, Ruest et al., 2003). It has not been implicated in innate immunity previously. Among POU transcription factor genes in *Drosophila, Dfr/Vvl, Pdm1/nub* and *Pdm2/miti* were identified in a screen for transcriptional regulators that bind the NF-κB-family transcription factor Dif; they are important for the control of AMP gene expression (Dantoft, Davis et al., 2013, Junell, Uvell et al., 2010, Junell, Uvell et al., 2007). The corresponding proteins were not, however, identified as physical interactors of Akirin in *Drosophila* (Bonnay et al., 2014). Thus if POU transcription factors do have a conserved role in regulating AMP gene expression, their precise function must have evolved, especially as nematodes lack Rel-family transcription factors (Pujol et al., 2001). Another transcription factor, LIN-40, a GATA protein, NuRD complex component and one of two *C. elegans* homologs of human metastasis-associated protein MTA1, was the top hit among AKIR-1’s binding partners. Recent genome-wide ChIP-seq data from the “model organism encyclopedia of regulatory networks” project (via www.encodeproject.org), revealed the presence of LIN-40 at the *nlp-29* promoter (binding site peak, V: 3984375) in DNA from uninfected young adult worms. This independent line of evidence supports the presence of a NuRD/AKIR-1 complex within this AMP gene cluster in the absence of infection. Consistent with our current understanding of its mechanism of action, we did not find STA-2 among the AKIR-1-interacting proteins. In the simplest model, a complex of AKIR-1, CEH-18 and the NuRD/MEC chromatin remodelling proteins is recruited to the *nlp* locus and opens it, but repress gene expression. Upon infection, the chromatin structure allows activated STA-2 access to the AMP gene promoters, and removal of the repressive NuRD/AKIR-1/CEH-18 complex permits gene expression. It is noteworthy that 3 of the 53 high-confidence AKIR-1 interactors are implicated in ubiquitin-mediated protein turn-over, and that *in vitro* ubiqutination activity can be detected within the purified NuRD/AKIR-1 protein complex specifically after infection, not before (JP, unpublished observations).

Chromatin remodelling at the promoters of immune genes can prime them for enhanced activation (Glass & Natoli, 2016). Many AMP genes in *C. elegans*, as in other species, are arranged in genomic clusters (Pujol et al., 2008b). AKIR-1-dependent modification of chromatin structure offers the possibility of coordinating a rapid increase in the expression of neighboring AMP genes, potentially important when faced with a fast-growing pathogen like *D. coniospora*.

Akirin functions together with the SWI/SNF complex in other species. Although we excluded a role for the *C. elegans* SWI/SNF complex in *nlp-29* expression, we did identify some SWI/SNF complex proteins, including SWSN-1, -3, -4 and -6, among the potential AKIR-1 binding partners. These were found through an unbiased whole-organism approach; it is likely that we sampled separate complexes from different tissues. Indeed AKIR-1 is known to be expressed widely and also to have essential functions in development; it is necessary for synaptonemal complex (SC) disassembly during meiosis (Clemons et al., 2013). These different candidates therefore merit investigation in the context of AKIR-1’s other functions. It would also clearly be of interest to attempt to recover AKIR-1 interactors specifically from the epidermis, but this is a technical feat beyond the current study.

SC disassembly involves a conserved RAS/ERK (Extracellular signal-regulated kinase) MAPK cascade. Interestingly, the same pathway is required for the response of *C. elegans* to infection by the Gram-positive bacterium *Microbacterium nematophilum* (Nicholas & Hodgkin, 2004). Within the rectal epithelium, it cooperates with a Gαq signaling pathway to trigger changes in cell morphology. At the same time, in motor neurons, Gαq functions independently of RAS signalling to influence nematode behaviour in the presence of *M. nematophilum* (McMullan, Anderson et al., 2012). Following infection, it also acts in the pharynx to regulate, non-cell autonomously, defence gene expression in the intestine (Gravato-Nobre, Vaz et al., 2016). These instances illustrate how the physiological response to infection is a mélange of interconnected signal transduction cascades. Further studies will be required to establish whether *akir-1* is required for any or all of these processes.

Across species, MAPKs act as regulators of chromatin structure. In yeast, the p38-related MAPK Hog1 physically interacts with the RSC chromatin-remodelling complex. This association is increased upon osmotic stress and is thought to direct the complex to bind osmo-responsive genes, changing nucleosome structure, increasing RNA polymerase II binding and causing a burst of transcription (Mas, de Nadal et al., 2009). In vertebrates, the SWI/SNF subunit BAF60 can be phosphorylated by p38 MAPK, also targeting it to specific loci (Simone, Forcales et al., 2004). It is not yet clear whether PMK-1 directly phosphorylates AKIR-1 or NuRD/MEC complex proteins; it was not found as a physical interactor of AKIR-1. The p38 MAPK PMK-3 on the other hand was. RNAi of *pmk-3* does not inhibit *nlp-29* expression (Zugasti et al., 2016). Notably, *pmk-3* does participate in adult axon regeneration, in a p38 pathway that while sharing some elements with the epidermal innate immune pathway (Kim, Thakur et al., 2016) is clearly distinct. Our results therefore raise the possibility that AKIR-1 plays a role in axon regeneration, in association with PMK-3.

We established that the SWI/SNF complex does not play a major part in modulating AMP gene expression in the epidermis. Rather the NuRD and MEC complexes, in a physical complex with AKIR-1 and CEH-18 play an essential role. One possible cause of this evolutionary re-wiring of a regulatory circuit could be the loss of NF-κB from nematodes, which has also led to a restructuring of the TLR pathway (Brandt & Ringstad, 2015). The precise evolutionary trajectories that led to these changes can only be the subject of speculation, but these lineage-specific adaptations likely reflect the extreme selective pressure that is exerted by pathogens. This plasticity is even more remarkable when one considers the essential developmental processes that many of these factors are involved in, limiting the degree of change that can be tolerated. In conclusion, as well as substantially advancing our understanding of immune defences in *C. elegans*, our results illustrate how an organism can evolve novel molecular mechanisms to fight infection while conserving an overall regulatory logic.

## Materials and Methods

### Nematode strains

All strains were maintained on nematode growth media (NGM) and fed with *E. coli* strain OP50 (Stiernagle, 2006). The wild-type reference strain is N2 Bristol. Strains carrying *akir-1(gk528), rde-1(ne300)* and the transgene [*ceh-18::TY1::GFP::3xFLAG*] (OP533) were obtained from the *Caenorhabditis* Genetics Center (CGC). Double mutants and strains containing multiple independent transgenes were generated by conventional crossing. The strains IG274 (containing *frIs7[nlp-29p::gfp, col-12p::DsRed] IV*) and IG1389 (containing *frIs7* and *frIs30[col-19p::GPA-12*,unc-53pB::gfp] I*) have been described elsewhere (Labed et al., 2012, Pujol et al., 2008a).

### Constructs and transgenic lines

Full genotypes of the transgenic strains are given below. The *akir-1p::AKIR-1::gfp* construct contains 1.6 kb of genomic sequence upstream of the start codon of E01A2.6 and was obtained by PCR fusion (Hobert, 2002) using primers JEP2091, JEP2092; JEP2108, JEP568, JEP569 and JEP570 and using genomic DNA and the vector pPD95.75 as templates. Microinjections were first performed using 20 ng/μl of the construct and the coinjection marker *myo-2p::mCherry* at a concentration of 80 ng/μl into N2 animals. Although transgenic strains were readily obtained, the observed fluorescence declined rapidly across successive generations (OZ unpublished observations). Since mutants in the Argonaute gene *rde-1* do not exhibit transcriptional silencing of transgenes in the soma (Grishok, Sinskey et al., 2005), we then performed the same microinjection but used *rde-1(ne300)* animals. From three independent transgenic lines generated, one was subsequently integrated using X rays and outcrossed three times with *rde-1(ne300)* generating *IG1550 rde-1(ne300) V; frIs32[akir-1p::AKIR-1::gfp; myo-2p::mCherry]*. This strain maintained transgene expression constantly across multiple generations. All additional strains carrying the *frIs32[akir-1p::AKIR-1::gfp; myo-2p::mCherry]* transgene were obtained by conventional crosses. The *akir-1p::gfp* construct was generated by PCR fusions using primers: JEP2091, JEP2092, JEP2095, JEP2096, JEP569 and JEP570 using genomic DNA, and the vector pPD95.75 as templates. Microinjections were performed using 20 ng/μl of the construct of interest and the co-injection marker *pNP135* (*unc-53pB1::DsRed*) at a concentration of 80 ng/μl in WT animals. Three independent lines were obtained and IG1485 was retained for further study. The single copy strain IG1654 carrying *AKIR-1::GFP* (*wt; frSi12[pNP157(akir-1p::AKIR-1::GFP)] II)* was obtained by CRISPR in N2 worms at the location of the *ttTi5605* Mos1 insertion (Vallin, Gallagher et al., 2012) and subsequent excision of the self-excising cassette (SEC) (Dickinson, Pani et al., 2015). pNP157 was made by Gibson cloning from a vector containing the SEC and recombination arms for *ttTi5605* (pAP087, kindly provided by Ari Pani) flanking *akir-1p::AKIR-1::GFP*, amplified from the strain IG1550, and the 3’UTR of *akir-1*, amplified from the wild type strain. The full locus *akir-1p::AKIR-1::GFP::3’UTR_akir-1* was confirmed by sequencing (primers available upon request). Microinjections were performed using pNP157 (*akir-1p::AKIR-1::GFP*) at 10ng/μl, pDD122 (sgRNA *ttTi5605)* at 40 ng/μl (kindly provided by Ari Pani), pCFJ90 *(myo-2p::mCherry)* at 2.5ng/μl, pCFJ104 *myo-3p::mCherry* at 5ng/μl and #46168 (*eft-3p::CAS9-SV40_NLS::tbb-2* 3′UTR; Addgene) at 30 ng/μl Roller worms that did not display red fluorescence were selected then heat shocked to remove the SEC by FloxP as described (Dickinson et al., 2015).

### Full genotypes of transgenic strains

IG274 *wt; frIs7[nlp-29p::gfp, col-12p::DsRed] IV* (Pujol et al., 2008b)

IG1389 *wt; frIs7 IV; frIs30[col-19p::GPA-12*,pNP21(unc-53pB::gfp)] I* (Labed et al., 2012)

IG1485 *wt; frEx547[akir-1p::gfp; unc-53p::DsRed]*

IG1502 *rde-1(ne219) V; Is[wrt-2p::RDE-1; myo-2p::mCherry]; frIs7 IV*

IG1550 *rde-1(ne300) V; frIs32[akir-1p::AKIR-1::gfp; myo-2p::mCherry]*

IG1555 *wt; frIs32[akir-1p::AKIR-1::gfp; myo-2p::mCherry]*

IG1575 *akir-1(gk528) I; rde-1(ne300) V; frIs32[akir-1p::AKIR-1::gfp; myo-2p::mCherry]*

IG1577 *akir-1(gk528) I; frIs32[akir-1p::AKIR-1::gfp; myo-2p::mCherry]*

IG1654 *wt; frSi12[pNP157(akir-1p::AKIR-1::GFP)] II*

IG1665 *wt; frSi12[pNP157(akir-1p::AKIR-1::GFP)] II; wgIs533[CEH-18::TY1::GFP::3xFLAG + unc-119(+)]*

### PCR fusion primers

The sequences of the primers used are:

JEP568: agcttgcatgcctgcaggtcgact,

JEP569: aagggcccgtacggccgactagtagg,

JEP570: ggaaacagttatgtttggtatattggg,

JEP2091: gatgaacaccgatagagagcaactg

JEP2092: gctctcgcggaaatgacgaat

JEP2095: agtgaaaagttcttctcctttactcattttacttctgaaagaaataatttgtggtta

JEP2096: atgagtaaaggagaagaacttttcact

JEP2108: agtcgacctgcaggcatgcaagctggagaggtacgaataggaatagtcat

### RNA interference

RNAi clones were from the Ahringer (Kamath, Fraser et al., 2003) and the Vidal (Rual, Ceron et al., 2004) RNAi libraries. Insert sequences were verified and target genes confirmed using Clone Mapper (Thakur, Pujol et al., 2014) before use. Epidermal-specific RNAi used the strain IG1502 *rde-1(ne219);Is[wrt-2p::RDE-1; myo-2p::mCherry];frIs7[nlp-29p::gfp, col-12p::DsRed]* (Zugasti et al., 2014). Worms were transferred onto RNAi plates at the L1 stage.

### Infection, wounding, osmotic stress and DHCA treatment

Infections, epidermal wounding and osmotic stress or dihydrocaffeic acid (DHCA) and treatments were performed as previously described (Zugasti et al., 2014).

### Killing and longevity assays

50–70 worms at the young adult stage were infected at 15°C for 1h with *D. coniospora* then transferred to fresh plates and the surviving worms were counted every day as described elsewhere (Powell & Ausubel, 2008). Longevity assays with *E. coli* strain OP50 used 50-70 worms at the young adult stage and were done at 20°C. The surviving worms were counted every day. Statistical analyses used one-sided log rank test within Prism (Graphpad software).

### Analyses with the Biosort worm sorter

Expression of *nlp-29p::gfp* reporter was quantified with the COPAS Biosort (Union Biometrica). Generally, a minimum of 80 synchronized worms were analyzed for size (TOF), extinction (EXT), green (GFP) and red (dsRed) fluorescence. The ratio Green/TOF was then calculated to normalize the fluorescence. When only mean values for ratios are presented, the values for the different samples within a single experiment are normalized so that the control worms (WT) had a ratio of 1. As discussed more extensively elsewhere (Pujol et al., 2008a), standard deviations are not an appropriate parameter and are not shown on figures with the Biosort. The results shown are representative of at least 3 independent experiments.

### RNA preparation and quantitative RT -PCR

RNA preparation and quantitative RT-PCR were done as described (Pujol et al., 2008b). Results were normalized to those of *act-1* and were analyzed by the cycling threshold method. Control and experimental conditions were tested in the same ‘run’. Each sample was normalized to its own *act-1* control to take into account age-specific changes in gene expression.

### qRT-PCR primers

Primers used for qRT-PCR are for:

*act-1:* JEP538 ccatcatgaagtgcgacattg JEP539 catggttgatggggcaagag;

*dcar-1:* JEP2030 cctacgctatttggtgcattggct JEP2031 tgcaccgaatcaccagaaacag;

*nlp-27:* JEP965 cggtggaatgccatatggtg JEP966 atcgaatttactttccccatcc;

*nlp-28:* JEP967 tatggaagaggttatggtgg JEP968 gctaatttgtctactttcccc;

*nlp-29:* JEP952 tatggaagaggatatggaggatatg JEP848 tccatgtatttactttccccatcc;

*nlp-30:* JEP948 tatggaagaggatatggtggatac JEP949 ctactttccccatccgtatcc;

*nlp-31:* JEP950 ggtggatatggaagaggttatggag JEP953 gtctatgcttttactttcccc;

*nlp-34:* JEP969 atatggataccgcccgtacg JEP970 ctattttccccatccgtatcc;

### Affinity co-purification assays

Affinity co-purification assays were performed as previously described (Chen, Cipriani et al., 2016) with minor modifications. From 3 independent mixed stage cultures of control *rde-1(ne300)* or *rde-1(ne300); akir-1(gk528)* worms carrying *akir-1p::AKIR-1::gfp,* samples were harvested, yielding about 4 g of flash-frozen pellets of *C. elegans*. In parallel, samples were also prepared from equivalent cultures that had been infected with *D. coniospora* for 16 h at 25 °C. Frozen samples were defrosted in a presence of lysis buffer (0.1% Nonidet P-40 Substitute, 50 mM Tris/HCl, pH 7.4, 100 mM KCl, 1 mM MgCl_2_, 1 mM EGTA pH 8.0, 10% glycerol, protease inhibitor cocktail (Roche), 1 mM DTT) and sonicated on Diagenode (cycle: 0.5 s, amplitude: 40-45%, 5 strokes/session, 10 sessions, interval between sessions: 30 s). After sonication, Nonidet P-40 Substitute was added up to 1% and the lysates were incubated with head over tail rotation at 4°C for 30 min, followed by centrifugation at 20,000 × g for 20 min at 4°C. Cleared lysate was then collected and split into either the anti-GFP agarose beads or the blocked control beads (40-50 μl, NanoTrap, Chromotek) (Fig. 5A). After head over tail rotation at 4°C for 60-90 min, the beads were washed once with lysis buffer containing 0.1% Nonidet P-40 Substitute, followed by two washings in each of the buffers I (25 mM Tris-HCl, pH 7.4, 300 mM NaCl, 1 mM MgCl_2_) then buffer II (1 mM Tris-HCl, pH 7.4, 150 mM NaCl, 1 mM MgCl_2_). Proteins were eluted by orbital shaking in 50 μl of glycine pH 2.6 followed by neutralization and ethanol precipitation. Precipitated proteins were resolubilized in 6 M urea/2 M thiourea buffer (10 mM HEPES, pH 8.0). Reduction and alkylation of proteins were then performed at room temperature, followed by digestion in solution sequentially using lysyl endopeptidase (Lys-C, Wako) for 3 h and trypsin (Promega) overnight as previously described (Paul, Hosp et al., 2011). Peptides were purified by solid phase extraction in C18 StageTips (Rappsilber, Ishihama et al., 2003).

### Immunoprecipitation assay

Mixed stage worms (IG1665) carrying AKIR-1::GFP and CEH-18::GFP::FLAG were harvested on ice and lysed in lysis buffer (0.5% Nonidet P-40 Substitute, 50 mM Tris/HCl, pH 7.4, 100 mM KCl, 1 mM MgCl2, 1 mM EGTA, 10% glycerol, protease and phosphatase inhibitor cocktail (Roche), 1 mM DTT), subjected to three cycles of freeze and thaw and sonicated on Diagenode (cycle: 0.5 s, amplitude: high, 5 min, interval between sessions: 30 s). Lysates were cleared by centrifugation. 200 μg of total protein was used for each immunoprecipitation: with anti-Flag (M2 clone, Sigma), and anti-HA as the unrelated control antibody (clone HA.11). Co-immunobound proteins were precipitated using Dynabeads Protein G matrix (ThermoFisher) and eluted in SDS buffer (1% SDS in TE, 150 mM NaCl). Immunoprecipitates were then resolved on a gel and subjected to Western blot analysis as described below.

### Liquid Chromatography Tandem Mass Spectrometry

Peptides were separated in an in-house packed analytical column (inner diameter: 75 μm; ReproSil-Pur C18-AQ 3-μm resin, Dr. Maisch GmbH) by online nanoflow reversed phase chromatography through an 8-50% gradient of acetonitrile with 0.1% formic acid (120 min). The eluted peptides were sprayed directly by electrospray ionization into a Q Exactive Plus Orbitrap mass spectrometer (Thermo Scientific). Mass spectrometry measurement was carried out in data-dependent acquisition mode using a top10 sensitive method with one full scan (resolution: 70,000, target value: 3 × 10^6^) followed by 10 fragmentation scans via higher energy collision dissociation (HCD; resolution: 35,000, target value: 5 × 10^5^, maximum injection time: 120 ms, isolation window: 4.0 m/z). Precursor ions of unassigned or +1 charge state were rejected for fragmentation scans. Dynamic exclusion time was set to 30 s.

### Mass Spectrometry Data Analysis

Raw data files were processed by MaxQuant software package (version 1.5.5.0) (Cox & Mann, 2008) using Andromeda search engine (Cox, Neuhauser et al., 2011). Spectral data were searched against a target-decoy database consisting of the forward and reverse sequences of WormPep release WS254 (28,071 entries), UniProt *E. coli* K-12 proteome release 2016_02 (4,314 entries) and a list of 245 common contaminants. Trypsin/P specificity was selected. Carbamidomethylation of cysteine was chosen as fixed modification. Oxidation of methionine and acetylation of the protein N-terminus were set as variable modifications. A maximum of 2 missed cleavages were allowed. The minimum peptide length was set to be 7 amino acids. At least one unique peptide was required for each protein group. False discovery rate (FDR) was set to 1% for both peptide and protein identifications.

Protein quantification was performed using the LFQ label-free quantification algorithm (Cox, Hein et al., 2014). Minimum LFQ ratio count was set to one. Both the unique and razor peptides were used for protein quantification. The “match between runs” option was used for transferring identifications between measurement runs allowing a maximal retention time window of 0.7 min. All raw mass spectrometry data have been deposited in the PRIDE repository with the dataset identifier PXD008074.

Statistical data analysis was performed using R statistical software. Only proteins quantified in at least two out of the three GFP pull-down replicates (or two out of two GFP pull-downs for the experiment using infected worms) were included in the analysis. LFQ intensities were log2-transformed. Imputation for missing values was performed for each pull-down replicate in Perseus (Tyanova, Temu et al., 2016) software (version 1.5.5.0) using a normal distribution to simulate low intensity values below the noise level (width = 0.3; shift = 1.8). The LFQ abundance ratio was then calculated for each protein between the GFP pull-downs and the controls. Significance of the enrichment was measured by an independent-sample Student’s *t* test assuming equal variances. Specific interaction partners were then determined in a volcano plot where a combined threshold (hyperbolic curve) was set based on a modified *t*-statistic (*t*(SAM, significance analysis of microarrays); *s_0_* = 1.5, *t_0_* = 0.9 ∼ 1.5) (Li, 2012, Tusher, Tibshirani et al., 2001). Proteins cross-reactive to the anti-GFP antibody were identified by a pull-down experiment using the non-transgenic *rde-1* strain and were filtered out from the AKIR-1 protein interactor dataset.

### Western blot analysis

Samples for western blot analysis were either prepared as per the co-precipitation protocol with the final elution performed in 50 μl 200 mM glycine pH 2.6 and immediately neutralisation by addition of 0.2 M Tris pH 10.4, or as per the immunoprecipitation protocol. Samples were then resolved on a 4-12 % BisTris Gel (Invitrogen) and subjected to transfer to a membrane.

Primary antibodies used in that study were as follow: anti-GFP (clone 11E5, Invitrogen, dilution 1:2000), anti-HDA-1 (Santa Cruz, dilution 1:2000), anti-LET-418 (kind gifts of F. Muller and C. Wicky, used at 1:500) and anti-FLAG (M2, Sigma, dilution 1:2000). The membrane was then incubated with horseradish peroxidase-conjugated secondary antibodies (1:10,000) at room temperature for 1 h, followed by brief incubation with substrates for enhanced chemiluminescence (Pierce ECL Plus).

### Chromatin immunoprecipitation

For extract preparations, N2 worms were grown on rich NGM seeded with HT115 bacteria, and young adult populations of worms were used to prepare about 3-4 gr of flash frozen worm popcorn. Worms were then fixed first with 1.5 mM EGS (ethylene glycol bis) for 20 min and then in 1.1% formaldehyde, with protease and phosphatase inhibitors, at room temperature with shaking, for 20 min. The fixing reaction was quenched by addition of glycine to a final concentration of 125 mM. Worms were then washed once with 10 ml FA buffer (50 mM HEPES/KOH (pH 7.5), 1 mM EDTA, 1% Triton X-100, 0.1% sodium deoxycholate, 150 mM NaCl) with protease inhibitors (Pierce), resuspended in FA buffer containing 0.1% sarkosyl and protease and phosphatase inhibitors, then dounce-homogenized on ice. Well-resuspended mixtures were then sonicated to shear chromatin (size rage 300-800 bp) using 12 cycles (30’ on, 30’ off) in a Bioruptor-Pico (Diagenode). Cellular debris was removed by centrifugation at 17,000 *g* for 15 min at 4 °C. Immunoprecipitation reactions contained approximately 3 mg of total protein, with 1% sarkosyl. Before addition of the antibody (NanoTrap-GFP, Chromotek), 5% of the material was taken as input. Immunocomplexes were collected and washed with 1 ml of the following buffers: FA buffer, two washes, 5 min each; FA buffer with 1 M NaCl, 5 min; FA with 500 mM NaCl, 10 min; TEL buffer (0.25 M LiCl, 1% NP-40, 1% sodium deoxycholate, 1 mM EDTA, 10 mM Tris-HCl, pH 8.0), 10 min, and TE (pH 8.0), two washes, 5 min each. Complexes were eluted in 1% SDS in TE with 250 mM NaCl at 65 °C for 30 min. Samples and inputs were treated with Proteinase K for 1 h, and cross-links were reversed at 65 °C overnight. DNA was purified with Qiagen PCR purification columns. Locus-specific ChIP qPCR reactions (SYBR Premix ExTaq II, TaKara) were done for each immunoprecipitation using specific elution (CHIP), negative control elution (nonspecific) and input samples, following a 50-fold dilution. Ct values were used to calculate the fold difference in DNA concentration between CHIP and nonspecific samples, normalized to the input.

*p_nlp-29:* JEP2521 gaaaaagaaacagagtctcgtgatg, JEP2527 tttctgattattaccacgtttttcg

*p_nlp-31:* JEP2529 cccagttcttcgtgtcaccac, JEP2530 gccgggcaaaatcacaaa

*p_nlp-34:* JEP2535 gacgtacctagacgtagaccatacacc, JEP2536 gtgacgtaattcgcaacatgg

*3’UTR_nlp-29:* JEP2544 ggggaagaaaataatttacatgagc, JEP2545 gcaagcgcaaaaatgttaaaaa

*3’UTR_nlp-31:* JEP2531 gcttttaataatatgacatgaccgaaa, JEP2532 gaaatttgacattcatcaaaatgct

*3’UTR_nlp-34:* JEP2539 ccgtacggatacggaggata, JEP2540 tttaaagtatattcgtcagcagcag

### Microscopy

Confocal images were captured using a confocal spinning disk (Yokogawa, Visitron Systems GmbH) microscope associated with a 512 x 512 pixels EM-CCD camera (Hamamatsu). Worms were immobilized in 0.01 % levamisole and visualized through a Nikon 40X oil, 1.3 NA objective and 1.5 lens, using a 488 nm laser.

## Acknowledgements

We thank F. Palladino and R. Waterston for discussion and sharing data before publication, Annie Boned for her contribution, F. Muller, A. Pani and C. Wicky for the generous gift of reagents. We acknowledge the PICSL imaging facility of the CIML (ImagImm), member of the national infrastructure France-BioImaging supported by the French National Research Agency (ANR-10-INBS-04). Worm sorting was performed using the facilities of the French National Functional Genomics platform, supported by the GIS IBiSA and Labex INFORM. This work was supported by institutional grants from INSERM, CNRS and Aix-Marseille University to the CIML and ANR program grants (ANR-12-BSV3-0001-01, ANR-16-CE15-0001-01, ANR-11-LABX-0054 (Labex INFORM) and ANR-11-IDEX-0001-02 (A*MIDEX)). Some nematode strains were provided by the CGC, which is funded by NIH Office of Research Infrastructure Programs (P40 OD010440)

## Author contributions

Conceived and designed the experiments: JP NP OZ JJE. Performed the experiments: JP JXC JS SO JB CT OZ. Analyzed the data: JP JXC OZ JJE. Contributed reagents/materials/analysis tools: JXC. Supervised the research: NP MS JJE. Wrote the paper: OZ JJE. All authors read and approved the final manuscript.

## APPENDIX

**Figure S1.**
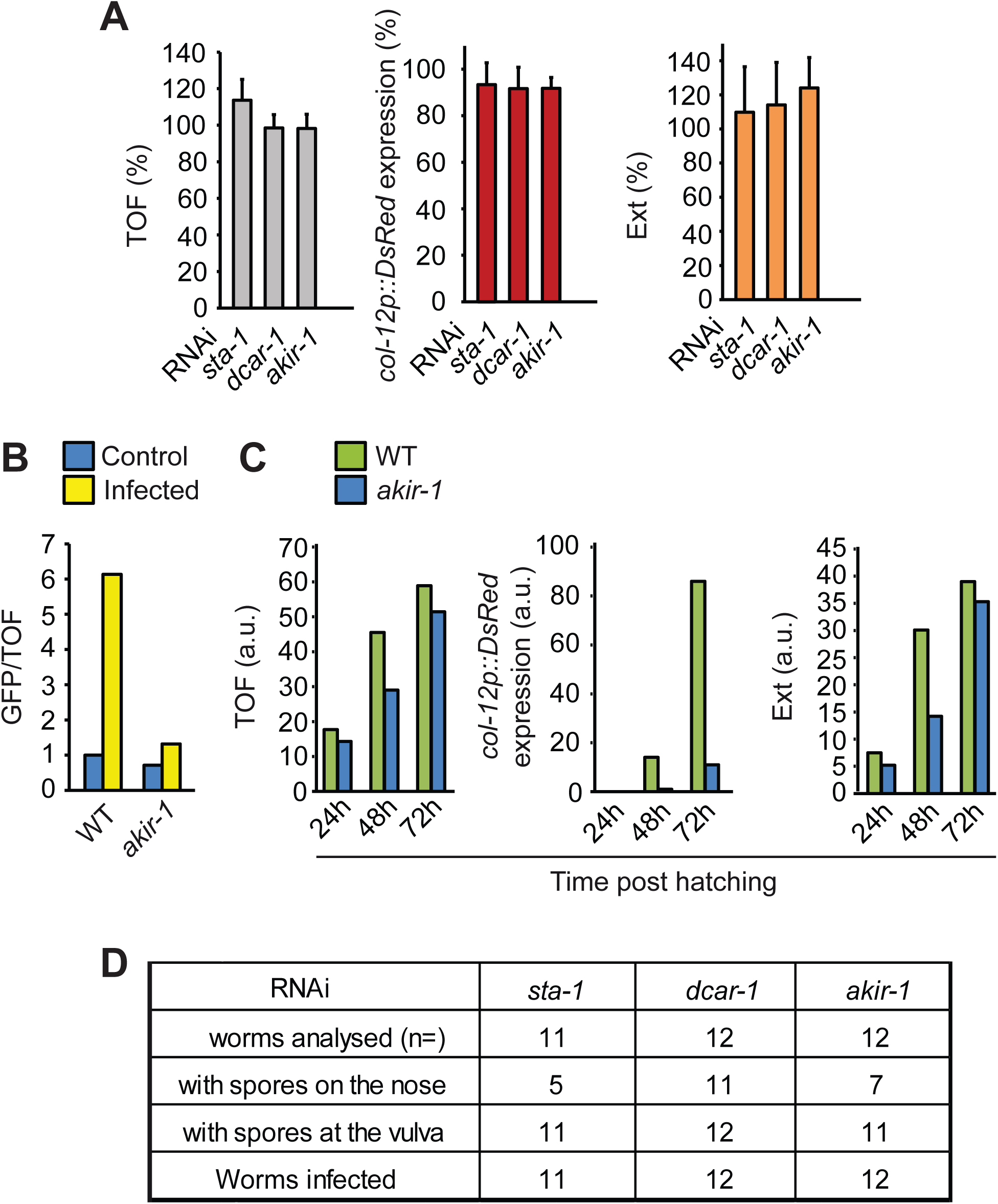
Akirin controls the expression of *nlp-29*, an AMP encoding gene. **A.** Quantification in arbitrary, but constant units of relative size (Time Of Flight; TOF; grey bars), *col-12p::dsRed* expression (red bars) and optical density (Extinction; Ext; orange bars) of wild type worms carrying the integrated array *frIs7* (which contains the fluorescent reporter transgenes *nlp-29p::gfp* and *col-12p::DsRed*) treated with RNAi against *sta-1, dcar-1* and *akir-1* and infected with *D. coniospora*. These results are taken from our previous genome-wide screen, from four independent experiments (average and SD). Values are relative to across-plate truncated means, as previously described (Zugasti et al., 2016). In all cases there are no significant differences between control and experimental values (paired two-sided student t test). **B.** Ratio of green fluorescence (GFP) to size (TOF) of wild type and *akir-1(gk528)* worms carrying *frIs7* and assessed without further treatment (control) or after infection by *D. coniospora*. Data are representative of three independent experiments. **C.** Comparisons of growth (left panel), *DsRed* expression (middle panel) and optical density (right panel) between wild type and *akir-1(gk528)* worms carrying *frIs7* on 3 successive days after hatching. Data are representative of three independent experiments. A minimum of 50 worms was analysed for each condition. **D.** Quantification of *D. coniospora* spore adhesion at the level of the nose and the vulva in wild type worms carrying the *frIs7* array and treated with RNAi against *sta-1, dcar-1* and *akir-1*.

**Figure S2.**
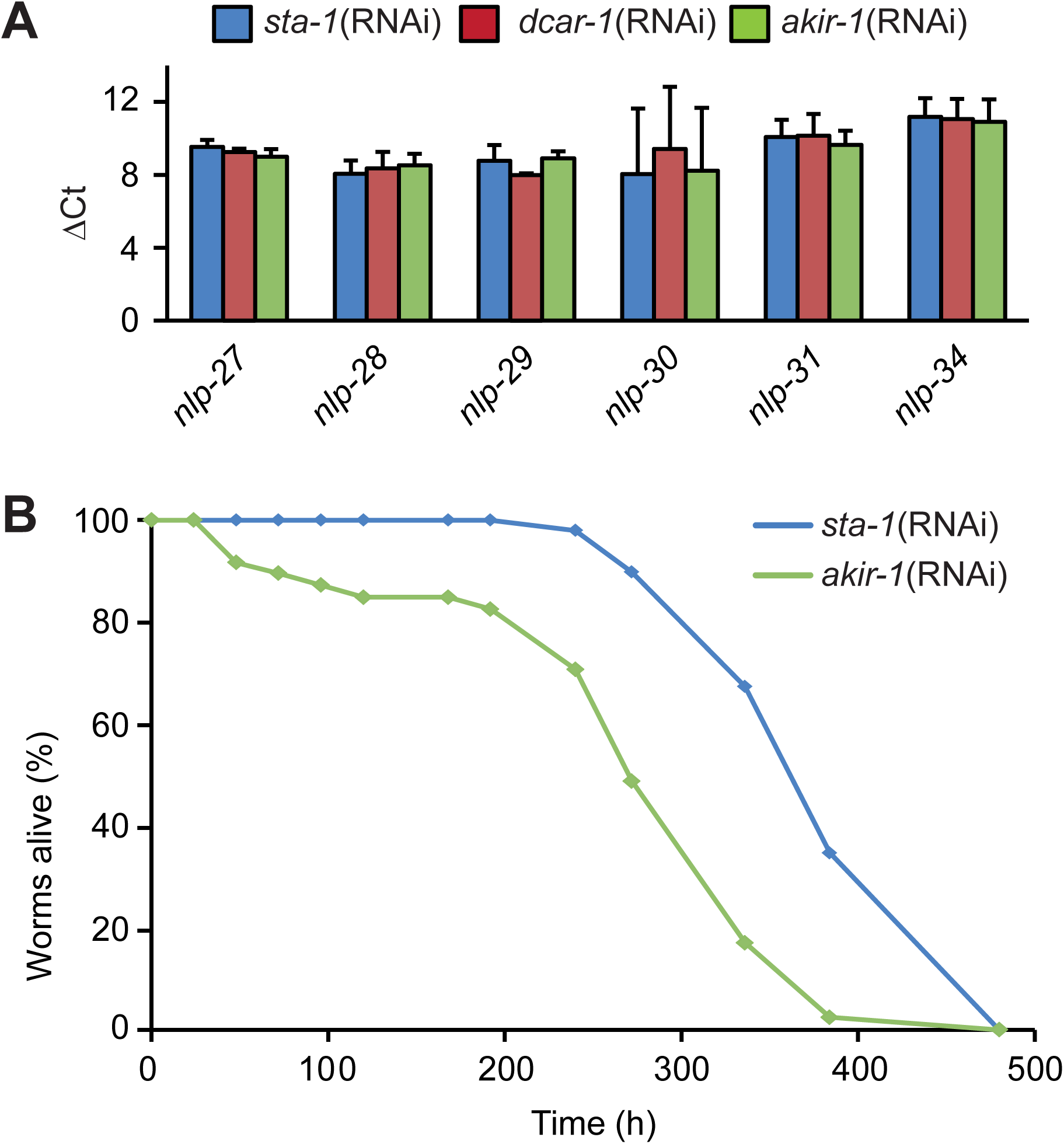
Akirin acts cell autonomously in the epidermis to control *nlp-29* expression and longevity. **A.** Abundance of mRNA for genes in the *nlp-29* cluster in *rde-1*(*ne219*); *wrt-2p::RDE-1* worms treated with RNAi against *sta-1, dcar-1* or *akir-1,* presented as the difference in cycling threshold (ΔCt) between each *nlp* gene and *act-1.* Data are from three independent experiments (average and SD). **B.** Survival of *rde-1*(*ne219*);*wrt-2p::RDE-1* worms treated with RNAi against *sta-1* (n=50) or *akir-1* (n=50). The difference between the *sta-1(RNAi)* and *akir-1(RNAi)* animals is highly significant (p*0.0001; one-sided log rank test). Data are representative of three independent experiments.

**Figure S3.**
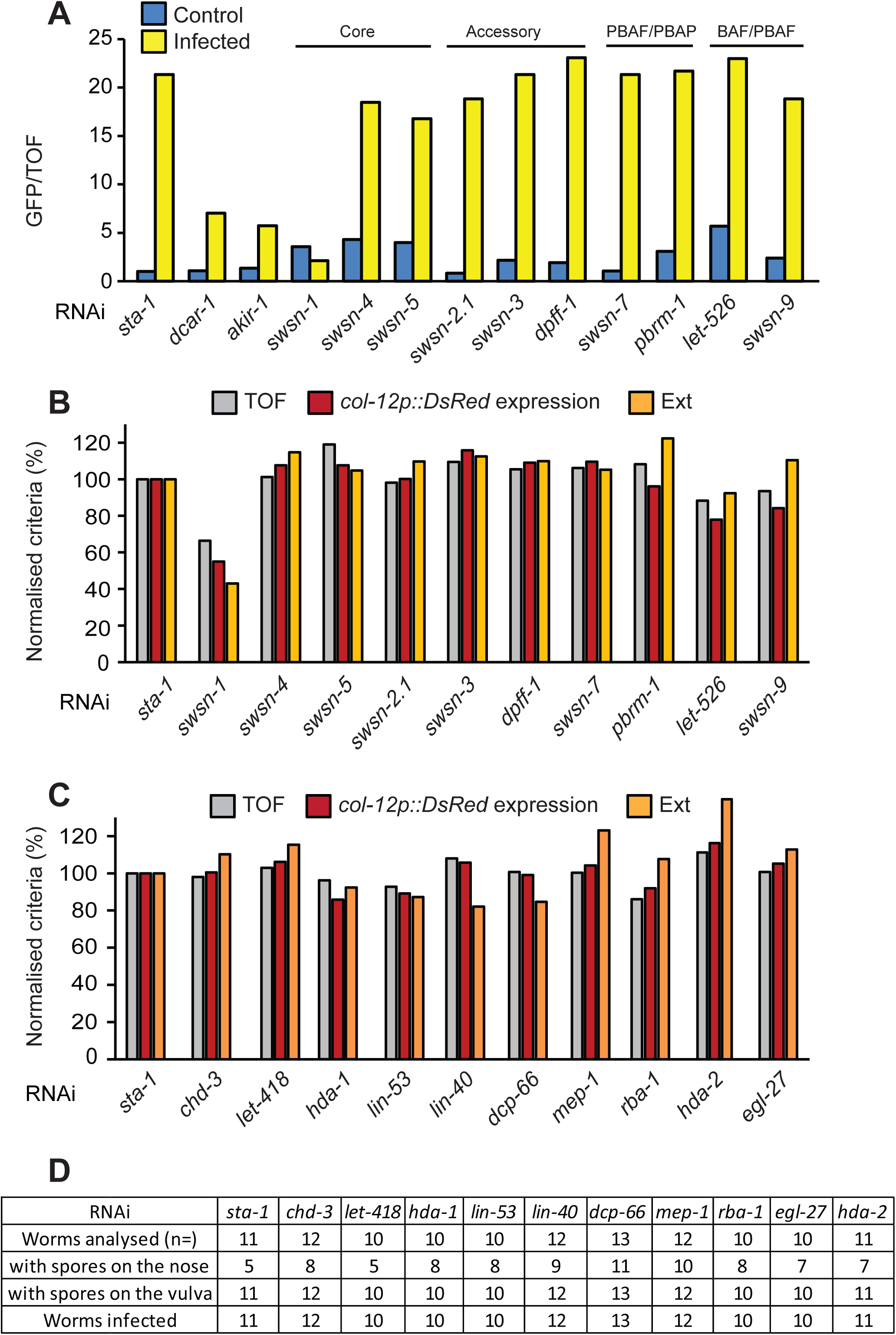
*nlp-29* expression is independent of the SWI/SNF nucleosome remodelling complex. Ratio of green fluorescence (GFP) to size (TOF) (A) and quantification in arbitrary, but constant units of relative size (TOF; grey bars), *col-12p::dsRed* expression (red bars) and optical density (Ext; orange bars) in wild-type worms carrying *frIs7* and treated with RNAi against different genes (**B, C**). The genes corresponding to core *(swsn-1, swsn-4, swsn-5),* accessory *(swsn-2.1, swsn-3, dpff-1*), PBAF/PBAP (*swsn-7 and pbrm-1*) and BAF/PBAF (*let-526 and swsn-9*) elements of the SWI/SNF nucleosome remodelling complex are indicated. Populations of >100 worms were analysed for each condition. Data are representative of three independent experiments. **D.** Quantification of *D. coniospora* spore adhesion at the level of the nose and the vulva in wild type worms carrying *frIs7* and treated with RNAi against the indicated genes.

**Table S1. Identification of protein-protein interactors for AKIR-1.** The quantitative results for analyses of 3 independent samples are given, referenced to Wormbase release WS254.

**Table S2. Comparison of protein-protein interactors for AKIR-1 and SWSN-2.2.** Data was from (Ertl et al., 2016) and Table S1. The annotations come from Wormbase (WS257). Gene identifiers were made uniform using Wormbase Converter (Engelmann, Griffon et al., 2011).

